# Nanomechanical Analysis of Extracellular Matrix and Cells in Multicellular Spheroids

**DOI:** 10.1101/193516

**Authors:** Varun Vyas, Melani Solomon, Gerard G.M. D’Souza, Bryan D. Huey

**Author notes:** **Mailing Address:** Department of Materials Science & Engineering 97 North Eagleville Road, Unit 3136 Storrs, CT 06269-3136 Office: IMS-158.

## Abstract

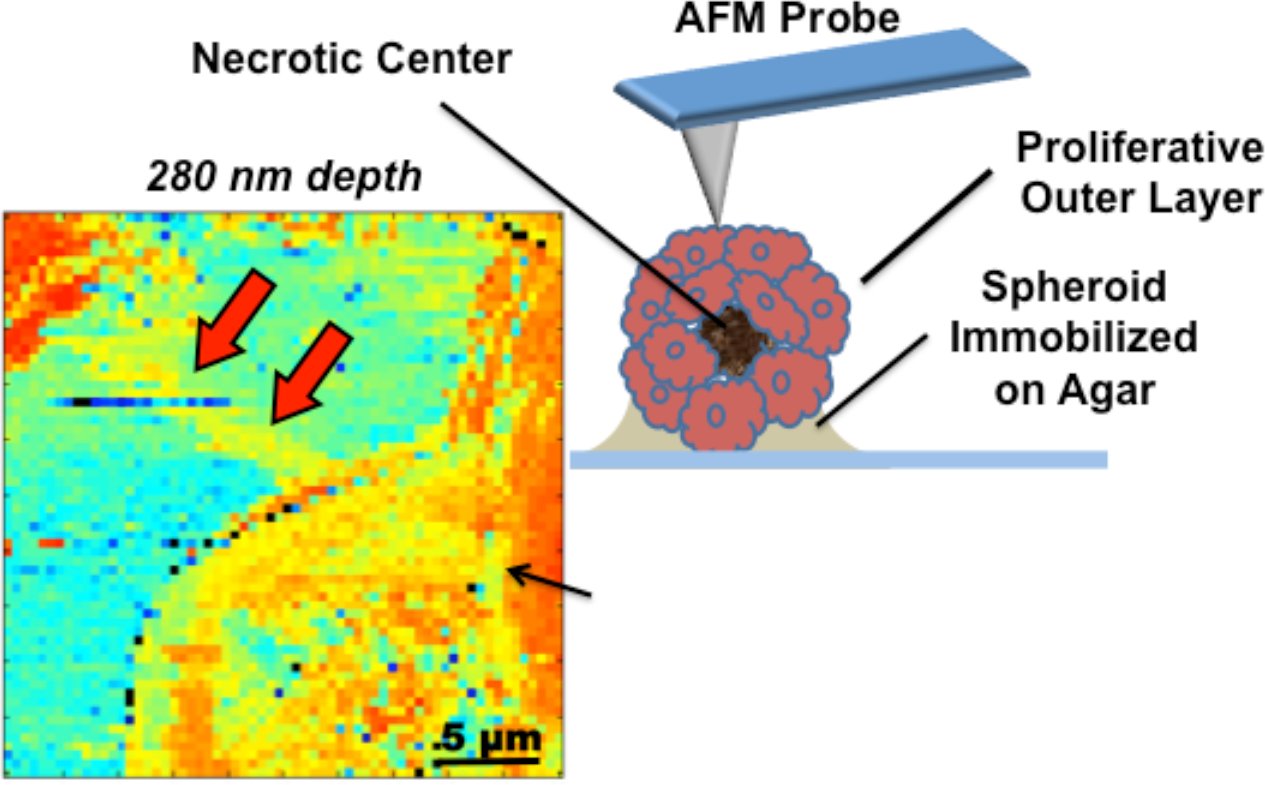

Nanomechanical investigation with Atomic Force Microscope has revealed new details regarding various nanomechanical heterogeneities for cells that are embedded in extracellular matrix in Multicellular spheroidal culture. This investigation sheds new insight into dynamic relationship of cells with their surrounding environment in tumors and 3D multicellular cultures.

Nanotechnology has revolutionized the field of cancer biology and has opened new avenues towards understanding nanomechanical variations in rapidly growing tumors. Over the last decade Atomic Force Microscope (AFM) has played an important role in understanding nanomechanical properties of various cancer cell lines. This study is focused on Lewis Lung Carcinoma Cell (LLC) tumors as 3D multicellular spheroid (MS). Such multicellular structures have enabled investigation of various components of tumors *in-vitro*. To better comprehend mechanical properties of cells and its surrounding extracellular matrix (ECM), depth dependent indentation measurements were conducted with Atomic Force Microscope (AFM). Force-vs-indentation curves were used to create stiffness profiles as function of depth. Here studies were focused on outer most layer i.e. proliferation zone of the spheroid. AFM investigations of sample a MS revealed three nanomechanical topographies, Type A- high modulus due to collagen stress fibers, Type B- high stiffness at cell membrane & ECM interface and Type C - increased modulus due to cell lying deep inside matrix at the a depth of 1.35 microns. Various nanomechanical heterogeneities revealed in this investigation can shed new light in developing correct dosage regime for various tumor dissolving drugs and designing more controlled artificial extracellular matrix systems for replicating tissue growth in-vitro.

**Short Statistical Summary:** This article describes nanomechanical characteristics of the cells embedded in extracellular matrix in a multicellular spheroid. The paper contains 6350 words including title page and references. Graphical Content contains 46 words. This article contains 6 Figures and zero tables.

## 1 Introduction

Tissues in higher living organisms have multilayered cell-cell and cell-extracellular matrix (ECM) interactions. Three dimensional in-vitro cell cultures used for investigating various kinds of tumors known as Multicellular Spheroids (MS), have become a preferred means to study and simulate an in-vivo environment^1^. 3D cell cultures with resembling structural and functional properties to real tissues has helped in understanding growth of tumorous, effects and penetration of toxins within a tissue and developing targeted drug-delivery systems^2–4^.

With the discovery of ECM in MS, gave us better insight towards developing 3D cell cultures^5^. Matrix act as both mechanical and biochemical shield MS that protects a cell. ECM also works as information gateway system that allows selective passage of signals via cell membrane. Here composition of ECM was found similar to that of tumors *in vivo*. Fibronectin, laminins, proteoglycans and collagen form major structural components within the matrix^5,6^. Structure forming elements like collagen and laminins play a major role in giving stability to MS.

MS being a tumor analog have inherent gradient of nutrients, oxygen and metabolites. In rapidly growing cultures, they have central necrotic core, which is surrounded by layer of quiescent viable cells and an outer most layer of rapidly proliferating cells. Central necrotic core and region of hypoxia is critical for testing anti-cancer therapeutics. Hypoxia have been identified as one of the cause of drug resistance in tumors^7–9^. Here penetration of drug through various layers of MS becomes a limiting factor towards drug efficacy. Therefore, this investigation is aimed towards understanding nanomechanical properties at surface of the spheroid where along with the cells, large volume of ECM is present that encapsulates multicellular structure. Various groups have extensively investigated mechanical characteristic of cancer cells, both metastatic and isolated cells from the tumors. But little know about their growth mechanics and impact ECM forming elements within a compact structure like a tumor or spheroid.

In this study, spheroidal cell cultures were developed using lewis lung carcinoma (LLC) cell line. LLC cell lines were derived from carcinoma of lung of a C57BL mice. LLC cells are well studied and are being used to determine outcome of drug induced toxicity by chemotherapeutic agents, developing methods to suppress metastasis, etc.^10,11^. There are number of methods like spinner flask, hanging drop, etc. that are being used to developing spheroidal cell cultures^1^. For the current investigation LLC cell spheroids were developed with liquid overlay method^12^. Understanding nanomechanical properties of spheroid via depth dependent indentation profiling (DDIP) of cells and surrounding ECM revealed minute changes in stiffness as a function of depth ^13^. Indentation dependent profiling methods have been explored by number of researcher in the field of AFM biology. Measurements conducted by *Rico et al* and similar groups primarily measured the modulus at one data point at time along the length of the force curve^14–16^. In this investigation, successive segments of Force-vs-distance curve were analyzed that provided ‘step-wise-modulus’ that helps in detecting variations in stiffness in 3-dimensions.

ECM is equally distributed within proliferation zone and quiescent zone. Hence the stiffness (i.e high density of ECM components) at the cell surface can be a controlling factor towards metastasis in tumors, drugs penetration and growth and proliferation of tumors. To map differential nanomechanics at the MS surface, force volume measurements were conducted with help of atomic force microscope (AFM). Other applications of AFM in the field biomaterials and cell biology include probing of biocompatible nature of metal oxide surface^17–19^, nanostethoscopy of cardiomyocytes and other living organisms^20–22^, Cell Nanomechanics^23–25^, etc.

In this communication our objectives is first, to highlight various nanomechanical heterogeneities within a MS and investigate interaction at cell-matrix interface. Second, to demonstrate application of multidimensional histograms along with the stiffness profiles in identification of such heterogeneities. Error parameters associated with local mechanical properties were also used in analysis. It helped in identifying small jitters experienced by AFM probe as it indents into a cell having well defined membrane as compared to ECM which is highly viscous mass of proteins, glycoproteins, etc. In our previous investigation with agarose gels, one could observe that stiffer gels had narrow error distribution when compared with agarose gels having lower stiffness^13^.

Earlier AFM measurements by Plodinec *et al* on rat spheroids were only focused on differential nanomechanical properties at the core and periphery of the tumorous structure^26^. This study is attempts to further explore nanomechanics of MS by mapping stiffness of cells and surrounding ECM. Surface scans were executed over 20 micron sized area with high resolution of 64 × 64 pixels, that brought forth three nanomechanical topographies into observation, Type -A i.e. stiffness of long range Collagen I stress fibers, Type- B i.e. high stiffness at interface of cell membrane & ECM, and Type-C i.e. indentation mapping up to depth 1.35 μm has also revealed stiffness of cells lying deep inside ECM. Modulus mapping at the surface was complemented with multidimensional histograms and depth wise error maps that gave crucial insight into mechanical diversity that constitutes an active 3D cell culture/ tumor in *in-vitro* environment.

## 2 Material and Methods

### 2.1 Production of spheroids

Lewis lung carcinoma (LLC) cells were obtained from ATCC (Manassas, VA). LLC cells were maintained in Dulbecco’s modified Eagle media (DMEM, Corning cellgros^®^, Manassas, VA) supplemented with 10% fetal bovine serum and 1% penicillin-streptomycin. Spheroids were produced using a liquid-overlay technique on agar surfaces. Briefly, 1% agar was coated in each well of a 96-well plate. Following the solidification of agar, 1000 LLC cells were plated in each well and the plate was centrifuged using a plate centrifuge (ThermoFisher Marathon 3000R) at 250xg for 15 min. The plate was placed in 5% CO_2_, 37°C atmosphere and the spheroids were allowed to grow for 6 days.

After 6 day of incubation spheroids were fixed on the glass coverslip using agarose gel. 0.1 % of warm agarose gel was allowed to cool down at room temperature and is gently poured over the coverslip. Before complete solidification of gel, a spheroid is carefully pipetted on its surface and lower half of the spherical mass of cell gets immobilized over the agar surface

### 2.2 Nanomechanical Measurements

AFM based force mapping measurements were conducted with an Asylum Research MFP-3D AFM with integrated inverted optics (Nikon TE-2000), in a liquid cell for complete immersion during the experiments. Arrays of 64 × 64 force indentations were acquired over a 20 μm × 20 μm area. All force maps were collected with indentation velocities of 32 μm/sec, amounting to individual indentations at a rate of ~2 Hz. Spring constants (0.3-1.3 nN/nm) of the Olympus TR800PB probes used throughout were calibrated following the widely applied Sader method^27^. Sneddon derived stiffness as a function of depth, as well as novel maps of the error in these calculations, were performed using custom analytical and visualization routines written with Matlab software.

## 3 Results

### 3.1 Experimental Setup

Spheroids were initially regarded as simple aggregates of cells with intimate cell-to-cell contacts. It was with discovery of ECM it became clear how closely they resembled tumors *in-vivo*(5). ECM is a gooey mass proteo-glycan, collagen and other structure forming elements. This membrane free extracellular component can have large variability in their mechanical physiognomies. This study is focused on such nanomechanical characteristics and of the cells embedded within this matrix forming a 3D aggregate of rapidly growing mass of cell (MS). LLC spheroids studied in this investigation were partially immobilized in agarose gel. The top view of cell fixed on the gel’s surface is show in Fig.1(a). Spheroid is 0.8 mm in diameter with a dark necrotic center marked by dotted red circle. Outer proliferative layer with lighter shades of grey is clearly visible with few loosely bound cells in its periphery. This provided stability to the spherical structure and prevented tumbling or any movement during long duration measurements via AFM probes (Fig. 1(b)).

**Fig. 1.**
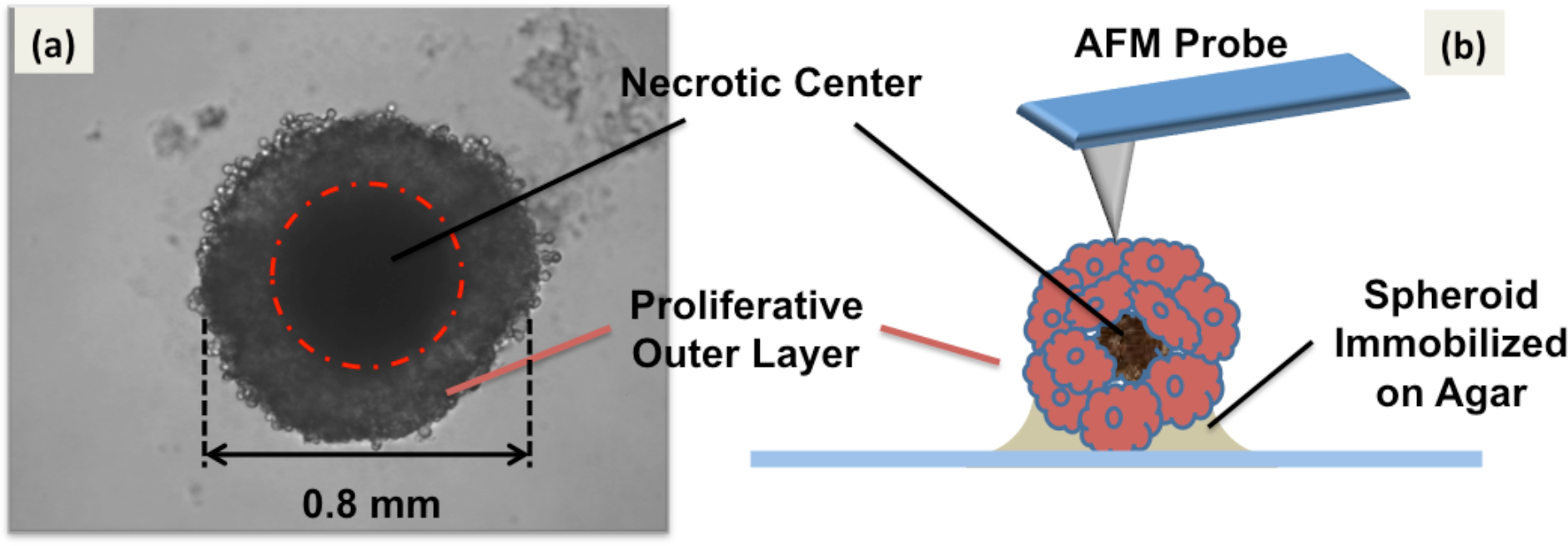
(a) Top view of a Spheroid partially embedded in agarose gel. Spheroid has diameter of 0.8 mm with a central necrotic core marked by dotted red circle. Outer proliferative layer having relatively lesser density of cells was used for nanomechanical investigations. (b). Schematic portraying experimental setup, lower half of the spheroid is immobilized in agarose gel. Upper half exposed to aqueous media is probed with AFM cantilevers.

AFM measurements were conducted over 20 micron-sized areas. 20 spheroids were examined to get most appropriate spot that presents both ECM and cells adjacent to each other. Each data set contains 4096 force-vs-indentation curves i.e. 64 × 64 pixels. Measurements were conducted at an indentation rate of ~ 2 Hz. It is a known fact that the measured mechanical stiffness depends on the loading rate^14,23^. This investigation required examining large number of samples, which in turn produced large volumes of data for analysis. Thus, in order to accelerate the experimental procedures and maintain viability of cells under the AFM probe, experiments were performed at a high indentation rate (~ 2 Hz). At lower rates, duration each scan becomes too long, this may kill or damage structural integrity MS under investigation. Therefore, the stiffness maps presented in this article, should be used to appreciate relative differences in stiffness between cells and ECM within a single data set.

### 3.2 Data Analysis

DDIP measurements were conducted using pyramidal AFM probe. Probe indents directly into viscous mass of ECM surrounding cells and at few places after indentation of few nanometers it comes in contact with cells embed in ECM. Due to square pyramidal geometry of the probe, modulus for developing indentation profile was calculated using Sneddon’s model.

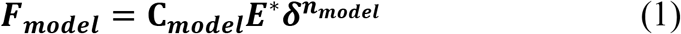

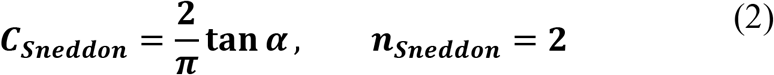

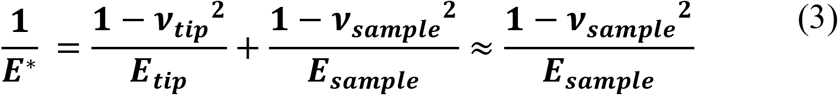

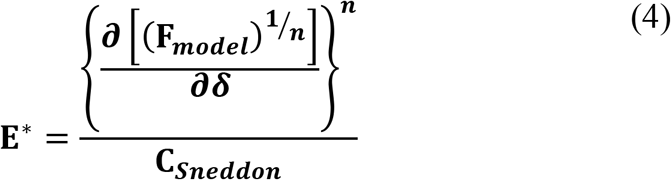

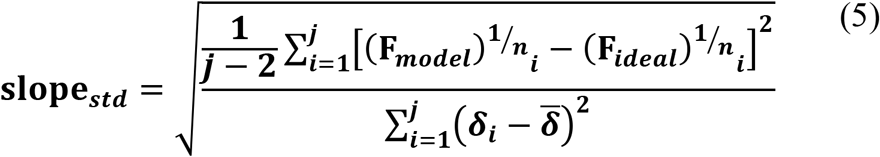

A simplified equation of measured force (***F*_*model*_**) vs indentation to a model-dependent power (***δ*^*n*^**) curves is shown as Equation-1. Other terms of the equation include a model dependent constant term (**C**_***model***_), and the local reduced modulus (**E\***). Equation for Sneddon’s mechanics and reduced modulus (E*) are presented as Equation 2 & 3 respectively. For calculating reduced modulus, Poisson ratio (***ν***) of 0.3 was employed for all samples. Modulus (E) of the tip is >100 GPa as compared to sample’s < 1MPa^28,29^. Hence, the first term in Equation 3 becomes negligible and is dropped from the final equation.

The local reduced modulus for any given Force-vs-indentation curve was calculated from slope of the n^th^ root of the curve, all is taken to the n^th^ power and is divided by the appropriate constant (Equation 4). This approach towards calculating the modulus also provides standard deviation of the modulus. Here standard deviation of the slope is calculated by simple linear regression in Equation 5 and is substituted by standard deviation of the actual slope in Equation 4(braces). Effective local modulus and its standard deviation at varying depths and at every pixel was calculated using Equation 3. Error maps developed here was used to interpret indentation data from novel perspective.

Data from each indentation curve is split into small fragments and each fragment is fitted with Sneddon’s model to estimate stiffness at certain depth is illustrated in Fig. 2a. All the force spectras from 64 × 64 indentations were plotted in a single multi-dimensional histogram (color contrast indicates histograms counts), shown as Fig. 1b. Due to the gelatinous nature of the cells and surrounding ECM repulsive forces rise gradually over a distance of 2-3 microns. Regions having relatively lower compliance, one can observe much slower increase in forces curves (0-10 nN/μm). To determine the point of contact with the sample, relatively high contact force of 4 nN was considered. At such high contact force there is sufficient deformation of cell around the AFM probe which has been predicted^30,31^ and observed^32^ by various research groups. This point of contact was latter used for developing topography of apical surface of the MS which otherwise cannot be observed using optical microscopy.

**Fig. 2.**
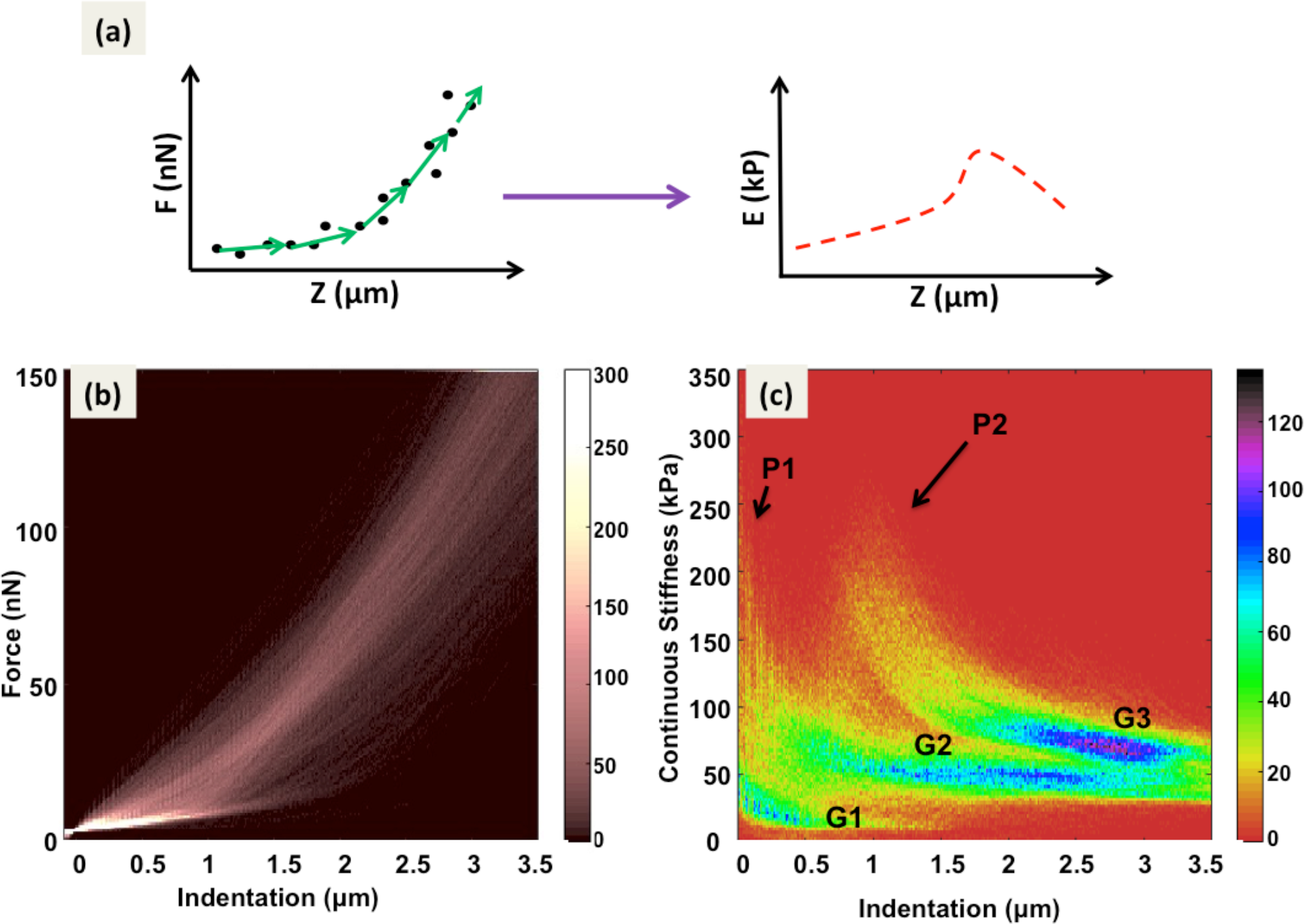
(a) Sketch shows stepwise fitting of Sneddon’s model into force curve data which gets converted into modulus vs. indentation curve. Modulus data from each pixel is plotted as stiffness map at various depths. (b) Multidimensional histogram (color contrast indicates the counts) of all 64 × 64 force (nN) versus distance (μm) curves in 20 × 20 μm scan of the surface of a spheroid (c) Multidimensional histogram of continuous stiffness (kPa) versus indentation (μm) of the force volume data shown in (b). Differential color contrast indicates density of the counts at different indentation depths in x axis, corresponding with the stiffness values in y axis.

As described in the illustration in Fig. 2(a) force-vs-distance (FD) curves were fitted with Sneddon’s model and continuous stiffness-vs-indentation multi-dimensional histograms were developed. A colored histogram (color contrast indicates histograms counts) with depth wise modulus is shown in Fig. 2(c). Plotting data from all continuous stiffness-vs-indentation curves in a multi-dimensional histogram, produces differential color contrast based on the density of counts at different depths (indentation). Here two peaks P1 and P2 are marked with black arrows. High density of counts creates three different groups with blue core (counts between 100-120) are labeled G1, G2 and G3. This contrast can only be appreciated with high resolution force volume measurements. Here 4096 curves were collected over a scan area of 20 μm, which gives a pixel size 312 nm. Such closely packed probing of the surface gives relatively higher number data points from the regions having similar nanomechanical characteristics. Therefore, heterogeneities in stiffness as a function of depth can be observed both from the histograms as well as from modulus profiles. Correlation between the histogram and modulus profiles is discussed latter in the text.

Surface topography of created from all the FD curve is shown in Fig. 3(a). Positioning of cells and ECM is explained with help of illustration in Fig.3(b). Bluish central region in Fig. 3(a) is ECM, which is illustrated with green color in Fig. 3(b). There are 3 cells (top left, right and bottom right) with light bluish green tone in topography (Fig.1 (a). The same are represented with red color in illustration in Fig. 3 (b). Here LLC cells seem to have spherical morphology. This was expected due their growth in tumor like environment and presence of large volume ECM in their immediate vicinity.

Data collected on measured stiffness as a function of depth (Fig. 2(c)) is plotted in form of a stiffness map shown in Fig. 3(c). Stiffness map presented here is at depth of ≤ 80 nm. ECM in the center of the Fig. 3(c) at ≤ 80 nm depth has large variations is modulus starting from 10 kPa and going upto 100 kPa. One can observe a large gradient in color contrast, from bluish green to red in color. To get maximum contrast, color bar for the modulus is presented in log scale. Presentation of the modulus in logarithmic scale can facilitates in identification of minute features having stiffness unlike rest of the matrix. During the measurement MS is suspended in growth media (DMEM). Continuous poking with the AFM probe can dissolve some of its matrix components into the surrounding aqueous environment. This may create large variations in stiffness at depths closer to the surface of the spheroid. ECM itself being gooey in nature, with mix various molecular and structural components can produce large variation in modulus observed close to the surface MS.

Cells earlier discussed in topography (Fig 3(a)) on top left, right and bottom right corner of the image have very high modulus when compare with adjacent ECM. Modulus of all the cells is > 100 kPa and goes upto 300 kPa. These high modulus values within first few 100 nm are also observed in multi-dimensional histogram and are labeled as peak P1 in Fig. 2(c). After first few 100 nm, there is a gradually decrease in stiffness as probe indents deeper into the cell. This decrease in modulus can be observed from stiffness profiles 280 nm, 800 nm, 1 μm and 1.35 μm in Fig. 4(a), 4(b), 4(c) & 4(d) respectively.

**Fig. 3.**
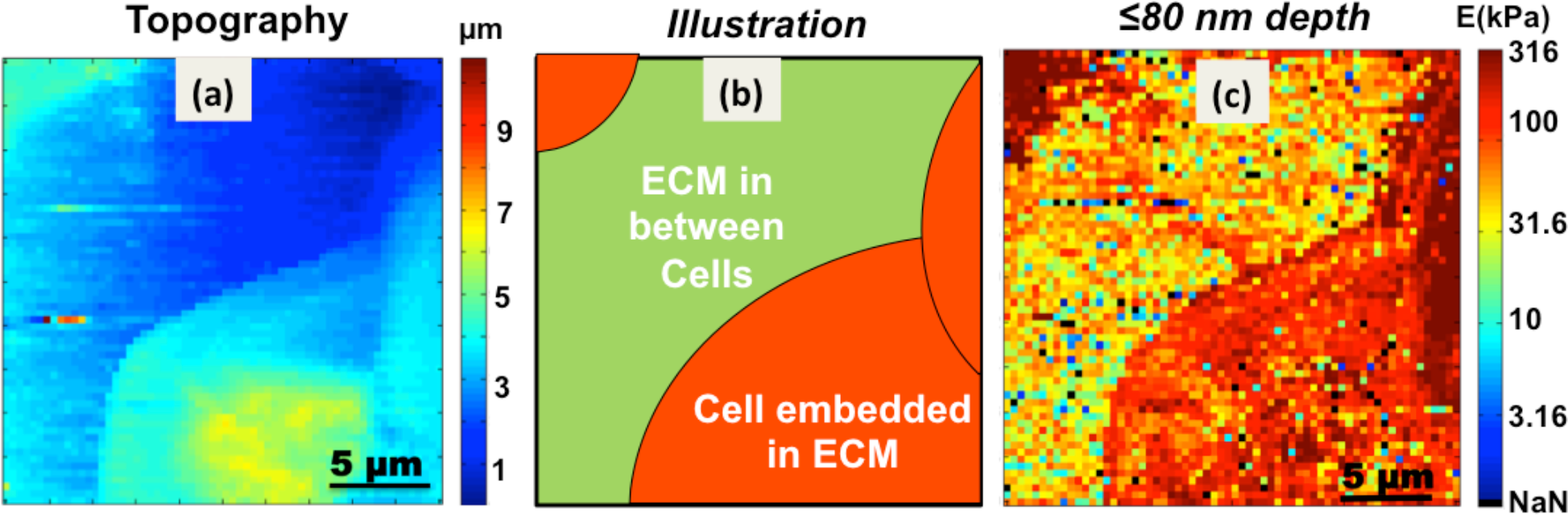
(a) Surface topography of LLC-spheroid cell created from 128 × 128 force curves. (b) Illustration explaining various nanomechanical features visible in stiffness profile. Large central green colored region is ECM in between cells and cells embedded in the matrix are colored red. (c) Indentation modulus profile of MS up to depth of 80 nm. Scan area of 20 × 20 μm having 64 × 64 pixels.

### 3.3 Sub-Surface Nanomechanics

Closer inspection of the maps derived from indentation depths of ≤ 80 nm to 1.35 μm revealed three types of nanomechanical topographies, Type-A, Type-B & Type-C. These features are explained by illustration in Fig. 5. The illustration highlights the point of contact of AFM probe with various sub-surface structural components (collagen - long elongated yellow fibers) in the cells (oval shaped) and surrounding ECM (green in color). Type-A nanomechanical features look like elongated fibrillary structures also known as stress fiber, can be observed in Fig. 4(a) (marked by two red arrows). They are possibly collagen based stress fibers that help cells to mechanically sense and respond to the presence of other cells over long distances (~ 100 μm). Collagen of types I, II III, V and XI are fibril-forming collagens ^33^. Studies show that when hMSCs are grown on type-1 collagen gels, they respond to stiffness of glass slide via 1000 μm of collagen gel ^34^. Similarly, when fibroblast are grown on fibrin gels, they can sense presence of other cells up to 250 μm away and are also known to respond by aligning themselves relative to each other ^35^. Therefore, stiffness and stability of MS can be directly correlated to modulus of these stress fibers. Collagen fibers are known to have modulus greater than 0.3 GPa but can have lower stiffness depending on the substrate and the source of the biomaterial^36–39^.

Stiffness of bulk ECM was measured up to few 10s of kilo Pascals in Fig. 4(a). Hence, the presence of type I collagen fibers should have stiffness greater than rest of biological material forming a MS. In Fig. 3(c), stress fiber observed in Fig. 4(a) looks like a smear in the center of the matrix. Its stiffness up to depth of 80 nm is ≥ 100 kPa. Collagen fibers have high stiffness but due to its presence close to surface of spheroid (top layer of ECM) it got masked by high degree of variability in modulus. At depth of 280 nm (Fig. 4(a)) it can be observed as an interconnecting fiber between cells separated by distance of 10 microns. Collagen fibers are known to have diameter greater than 250 nm and sometimes they might be as thin as 30 nm ^40,41^. Accurate estimation of modulus of stress fiber in MS will also depend on the loading rate, plus fibers are spread across a squishy matrix of biomolecules. Fibers will get pulled and pushed around as probe indents into the spheroids. Hence, stress and strain factors will also come into play for more precise measurement of modulus of collagen fibers in 3D cell cultures^42^.

**Fig. 4.**
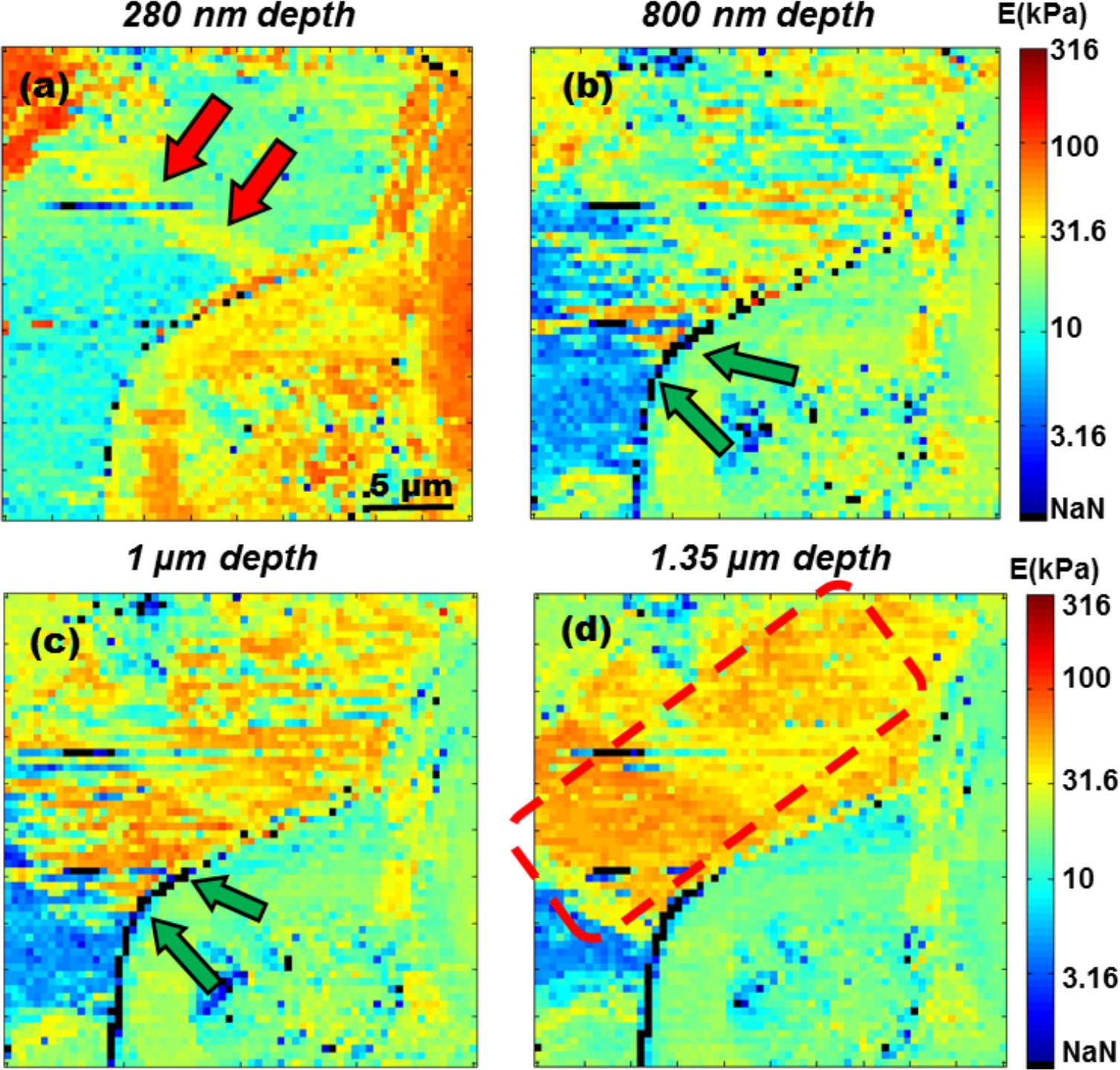
Stiffness maps of MS at various depths (a) At depth of 280 nm, modulus of ECM is at 10 kPa and cells have stiffness < 100 kPa. Double red arrows mark interconnecting collagen type I stress fiber. (b) Interface of cell membrane and ECM at depth of 800 nm is marked by double green arrows, has stiffness greater than 316 kPa, hence observed as string of black pixels. (c) Double green arrows mark regions of high stiffness at cell membrane-ECM interface at 1 μm depth. (d) At 1.35 μm depth, cell embedded deep inside the ECM can be observed with stiffness > 31.6 kPa is marked by dotted red square.

Type-B nanomechanical features illustrated in Fig. 5, arise due to interactions taking place between ECM and cell membranes. When cells are grown as monolayered cultures, apical surface is exposed to aqueous media whereas *in-vivo* most cells are embedded in ECM. Cells associated with fluid connective tissues like blood and lymph can be found floating or exposed to aqueous environment. There is hardly any interaction with any viscous mass like that of ECM other than their immediate aqueous environment. Therefore, stiffness of a cell might be influenced by interaction between the cell membrane and surrounding extracellular matrix. In this investigation it was interesting to observe that where cell membrane was in direct contact with ECM, a very high modulus ≥ 300 kPa was observed. As the probe indents into membrane-matrix interface (MMI) a sharp increase in stiffness was observed. Regions of high stiffness at MMI are marked by green arrows in Fig. 4(b) & 4(c) at depth of 800 nm & 1 μm respectively.

Here pixels having modulus greater than 316 kPa are shown with a black contrast, labeled as NaN in color bar. For the sake of visual presentation of sub-surface nanomechanical topographies and to accommodate maximum number nanomechanical heterogeneities logarithmic scale was used between 1kPa to 316 kPa. Stiffness for most of the soft biological material falls within this range. Hence, modulus greater than the cut-off value of 316 kPa is denoted by black contrast pixels and are shown in color bar below 1 kPa, hence labeled NaN.

Cell present on the outer most later of MS also have thin layer of ECM over them. As AFM probe indents into the surface, cell with a coating of ECM should have higher stiffness as compared to rest of the cell body. This effect was earlier observed in Fig. 3(c) (depth ≤ 80 nm) where cells on the lower right and top left corner have modulus up to 300 kPa. Hence, the cells in top layer of MS also fall into Type-B category. This explains why there is significantly higher stiffness up to indentations of 100 nm. Thereafter, gradual decrease in modulus was observed, as AFM probe has indented deeper into cytoplasm of the cell, that has low stiffness due to is gelatinous nature.

This increase in stiffness at MMI can be attributed to formation of basal lamina assembly at the cell surface^43^. ECM architecture near the cell surface is composed of interconnected polymers of Collagen IV and VII^44^. They are bound to laminin sheets, which are organized at the cell membrane by integrins and dystroglycan. Binding of collagen IV network to laminin takes place via nidogen molecules^45^. Assembly is further strengthen by binding to Collagen VII dimers^6,46,47^. A conclave of structural proteins over the cell membrane strengthens basal lamina. The combined effect of these tightly packed structures can considerably increases modulus of the cell membrane which may go above 300 kPa. Therefore, cells embedded in ECM in spheroids have shell like structure around them that provides them with lot of mechanical stability.

**Fig. 5.**
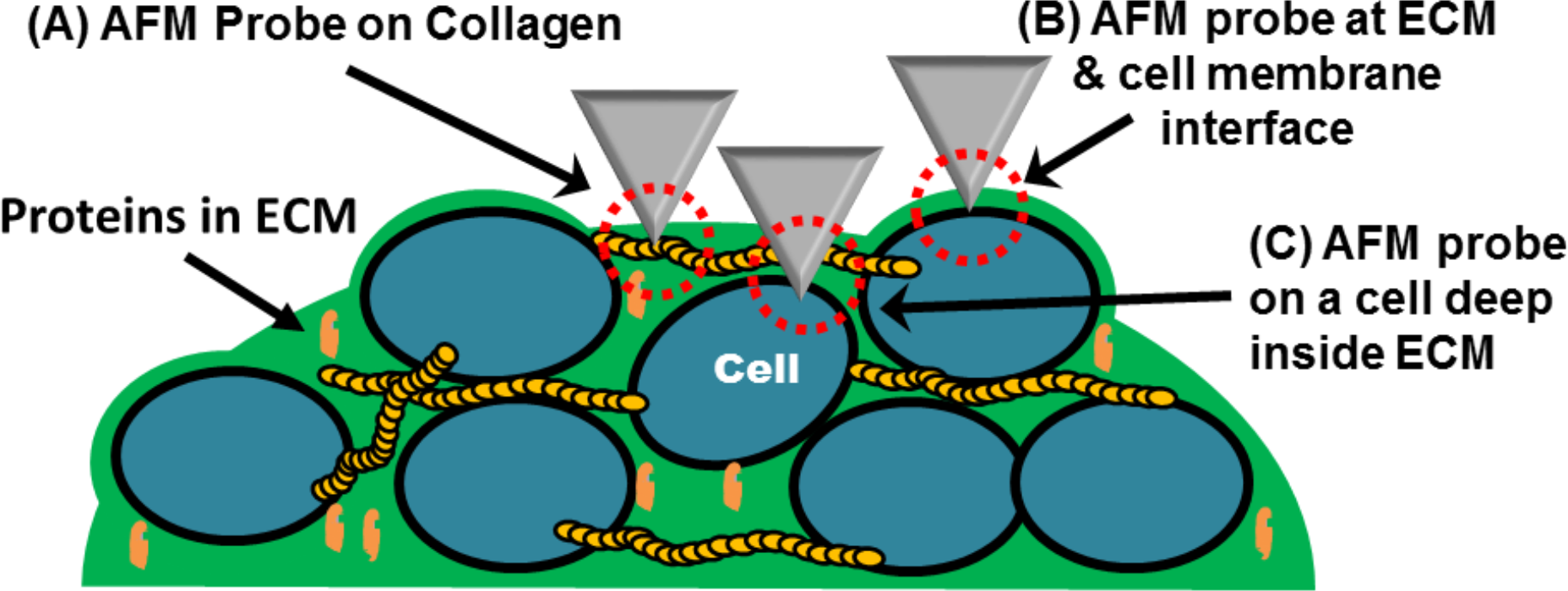
Sketch describes three types of nanomechanical topographies identified with DDIP. Type (A) is long range collagen type I stress fibers. They have considerable high stiffness as compared with surrounding ECM. Type (B) is region of high stiffness at interface of cell membrane and ECM. Type (C) is a cell embedded deep inside the ECM. They are identified by gradual increase of stiffness as AFM probe indents deeper into the ECM.

The last nanomechanical topography is Type-C, which is illustrated in Fig. 5 and is marked by red square in Fig. 4(d). Outer surface of spheroid doesn’t have an even distribution of cells. There are gaps filled with viscous ECM. As probe indents into the regions filled with ECM there is a decrease in stiffness from the depth of ≤ 80 nm (Fig. 1b) to 280 nm (Fig. 2a). Average stiffness of the ECM goes down from 31.6 kPa to 10 kPa. Only collagen fiber marked by double red arrows has relatively higher modulus as compared to the surrounding ECM. Furthermore at indentation depths of 800 nm (Fig. 4(b)), 1 μm (Fig. 4(c)) and 1.35 μm (Fig. 4(d)), one can observe a gradual increase in stiffness that goes up to 100 kPa. As illustrated in Fig. 5, Type-C could be due to a cell underlying deep inside the ECM. This effect can also be observed in multi-dimensional histograms. This second rise in modulus, labeled as Peak P2 can be observed in histogram in Fig. (c). This increase of modulus is from 0.5 μm (indentation) onwards and considerably number of counts having stiffness greater than 100 kPa can be observed up to depths of 1.5 μm. This observation of Type C features is limited by length of the AFM probe. Olympus TR800PB probes have height of 3 ± 0.5 μm, hence the analysis of the histogram data is considered up to depth of 3.5 microns. ECM being highly viscous, it tends to adhere to AFM probe. Any further investigating of nanomechanical properties of cells in quiescent zone of the spheroid will require development of specialized AFM probes.

Referring back to the histogram in Fig. 2(c), grouping of the counts can also be correlated with number pixels with similar values of stiffness. Group G1 is primarily contribution of modulus values from ECM up to depths of 500 nm. As the probe indents deeper, modulus averages out at all depths hence the grouping of counts is labeled as G2. Highest density of counts is observed beyond 1.5 μm. Downward trend from peak P2 merges into group G3. This third grouping of modulus is observed from indentations between 2 μm to 3 μm. Here higher density of counts can be due to Type-C nanomechanical topographies. They have relatively higher average stiffness of 100 kPa. Cells that are deep inside the spheroid are surrounded by ECM and other cells in their vicinity. This would put more mechanical load on to the cells present in the sub-surface regions. Hence, they might have higher modulus as compared cells that are present in the outer periphery.

### 3.4 Indentation Error Maps

Standard deviation from slopes of each FD curve was used for developing error profile at different depths. Error profiles at depths of ≤ 80 nm, 500 nm and 1 μm are shown is Fig. 6. Error profiles are correlated with previously discussed stiffness maps at different indentations. At ≤ 80 nm (Fig. 6(a)), ECM has higher error values as compared adjacent cells in top left, right and lower right corner of the image. For ECM, observed standard deviation of the calculated modulus (percentage error) was greater than 1. This might be due to lack of clearly defined structures or membrane on the outer most layer spheroid. Gooey nature of the ECM will also have significant contribution towards higher percentile error, especially in regions or spaces between cells filled with ECM.

**Fig. 6.**
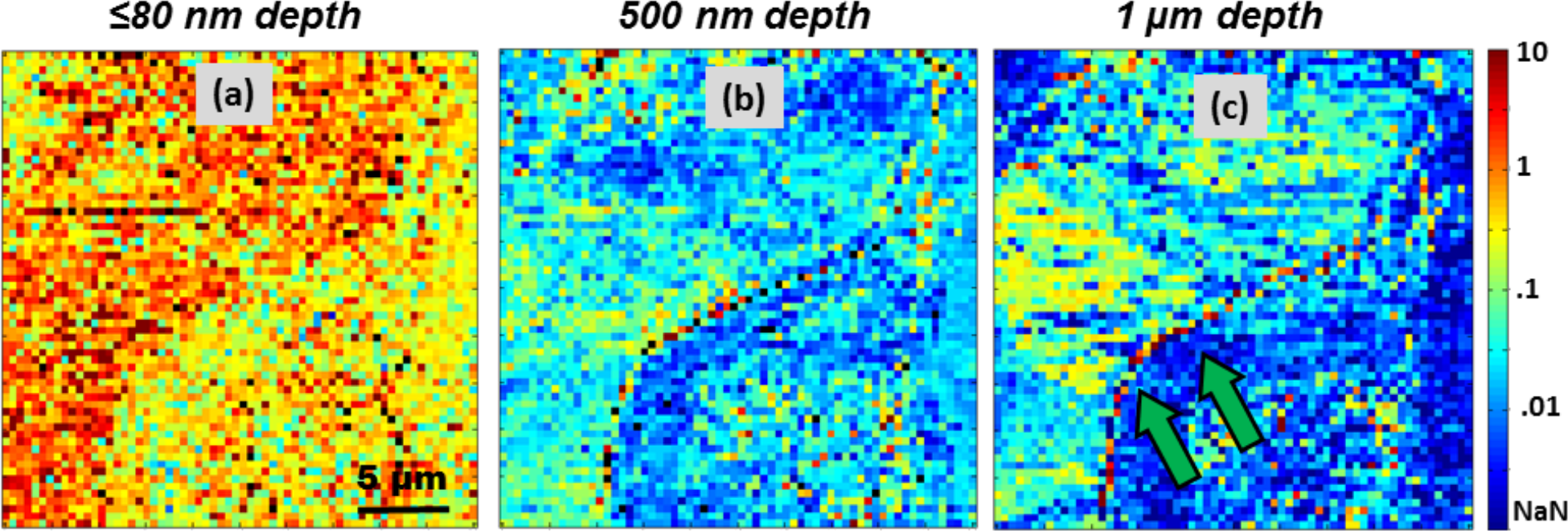
Standard deviation of calculated modulus is shown as percentage error maps at various depths (a) Error profile of stiffness up to depth of 80 nm. ECM has higher percentile error as compared to cells forming MS. (b) At 500 nm depth, error is < 1 for both cells and ECM. (c) Probe indents deeper to a depth of 1 μm. Edge of the cell interacting with ECM has higher error. Region of cell membrane - ECM interface is marked by two green arrows has percentile error > 1.

On closer examination of error profile in Fig. 6(a) & 6(b), i.e. from ≤ 80 nm to 500 nm there is a decrease in error to 0.01 units. At 1 micron depth Fig. 6(c) regions with higher modulus have errors greater than 0.1 unit. At depth of 1 μm, MMI regions have higher error values that go up to 10 units and a reddish parabolic streak across the edge of the cell is marked by two green arrows. This high error could be the result of resistance experienced by the probe as it indents deeper around the edges of the cell. Stiffness at MMI is greater that 300 kPa, when probe penetrates through the interface it can experience lots of jitters, thereby introducing relatively more errors. These errors are mildly visible in error map at 500 nm depth and significantly masked at ≤ 80 nm due to the squishiness of the ECM.

## 4 Discussion

Main objective of culturing MS is to replicate tumorous environment *in-vitro* for developing therapeutics that can penetrate and kill any cancerous growth inside the human body. Due to layered structuring and compactness of tumors, breaching outter most layers of cells including targeted drug delivery of drug becomes a limiting factor towards curing a cancer patient. ECM in tumors is known to present transport barriers that restrict drug penetration inside a solid tumor. Treatment of collagenase prior to delivery of 100 nm sized nanoparticles has found to increase particle penetration by 4-fold^48^. Both Type A & B topographies are collagen based. Collagenase treatment when coupled with nanomechanical studies can help in building correct dosage regime. Understanding tumor micro environment can therefore play a crucial role in developing effective anticancer drugs^4^. By measuring its stiffness one can look for better means to dissolve outer mass of cells and develop methods to deliver therapeutics into the quiescent zone for complete dissolution of the multicellular structure^2^.

Composition of ECM in MS is considerably different when compared with monolayer cultures ^49^. Previous studies have also suggested increased stiffness of ECM in malignant phenotypes/cultures and is associated with increased concentration of type I collagen in the matrix^50–52^. Plus type IV collagen being most abundant constituent of basement membrane, is associated with tumor fibrosis and accumulates in tumor interstitium ^46,53,54^. Therefore, detection of high stiffness at MMI by AFM probe is consistent with studies conducted on ECM and tumorous tissues so far.

Another interesting application of depth dependent investigations would be to investigate of artificial ECM. With DDIP, scanning surfaces of the cultures with artificial ECM would help fine tune nanomechanical properties most appropriate for growing 3D cell cultures. Some groups have started to develop permissive hydrogels that mimic ECM in MS. They have substituted protein based ECM components with poly(ethylene glycol) (PEG)^55–57^. Such artificial matrices can only act as a physical scaffold to hold cells proliferating in a 3-dimension (3D). But the exact nanomechanical properties in a native ECM environment are controlled by collagen, laminins and other components. Tissues engineered with artificial ECM can have different mechanism towards delivery of therapeutics and nutrients in 3D cultures. Therefore, information gather via nanomechanical investigation of matrix and cells in a MS can help in improving hydrogel based replacements for ECM.

The main limitation in conducting force measurements over spheroids is its opaque nature. Significant number of hit & trial are required to find optimum region in its outer periphery. Hence, it was not possible to conduct simultaneous fluorescence or optical measurements. But with the development of Light-sheet-based fluorescence microscopy (LSFM)^58^, one day it might be possible to combine force measurements with LSFM that would give us fine control to probe and selective study mechanical properties of cells in proliferation and quiescent zone in MS.

## 5 Conclusion

Some groups have reported extensively analysis of force curves to determine stiffness of living tissues but most of the investigations are focused on measuring modulus to a depth of few nanometers. Here we report for the first time how depth dependent indentation profiling (DDIP) can gives us crucial insight into nanomechanical diversity in a complex agglomerate of cells and extra-cellular matrix. Other investigators have conducted similar studies but they have worked with spherical probes having diameter greater than 5 micron^28^. There area of contact with cells grown on flat substrate would be much greater 312 nm. Even with pyramidal probes resolution was not more 16 × 16 pixels. Due to poor resolution and relatively greater area of contact with spherical probe depth dependent investigations won’t reveal any significant sub-surface heterogeneities. This study puts emphasis on minute changes in stiffness that are result of interaction between various components present within rapidly proliferating mass of cells. This novel approach of stiffness profiling can have profound application in 21^st^ century as we move towards synthetic bio-mechanical platforms. Knowledge regarding dynamics of interactions of cells with their surrounding environment is essential for developing any kind artificial 3D cell cultures. The same can also give insight towards targeting tumorous tissues i.e. mechanical stiffness at MMI can significant impact on bio-availability of the drug towards the treatment of concerned biological disorder.

## Author Contributions

All authors have given approval to the final version of the manuscript.

## Notes

Authors declare no competing financial interests.

## References

1 R.-Z. Lin and H.-Y. Chang, Biotechnol. J., 2008, 3, 1172–1184.

2 G. Mehta, A. Y. Hsiao, M. Ingram, G. D. Luker and S. Takayama, J. Control. Release, 2012, 164, 192–204.

3 S. H. Jang, M. G. Wientjes, D. Lu and J. L.-S. Au, Pharm. Res., 2003, 20, 1337–1350.

4 A. I. Minchinton and I. F. Tannock, Nat Rev Cancer, 2006, 6, 583–592.

5 T. Nederman, B. Norling, B. Glimelius, J. Carlsson and U. Brunk, Cancer Res., 1984, 44, 3090 LP–3097.

6 J. K. Mouw, G. Ou and V. M. Weaver, Nat. Rev. Mol. Cell Biol., 2014, 15, 771–785.

7 S.-H. Kim and H.-J. K. and C. R. Dass, Curr. Drug Discov. Technol., 2011, 8, 102–106.

8 F. Andre, N. Berrada and C. Desmedt, Curr. Opin. Oncol., 2010, 22, 547–551.

9 M. PV Shekhar, Curr. Cancer Drug Targets, 2011, 11, 613–623.

10 J. S. Bertram and P. Janik, Cancer Lett., 2017, 11, 63–73.

11 M. S. O’Reilly, L. Holmgren, Y. Shing, C. Chen, R. A. Rosenthal, M. Moses, W. S. Lane, Y. Cao, E. H. Sage and J. Folkman, Cell, DOI:10.1016/0092-8674(94)90200-3.

12 J. Carlsson and J. M. Yuhas, Recent Results Cancer Res., DOI:10.1007/978-3-642-82340-4_1.

13 V. Vyas, M. Solomon, G. G. M. D’Souza and B. D. Huey, J. Nanosci. Nanotechnol., 2018, 18, 1557–1567.

14 Y. M. Efremov, A. A. Dokrunova, D. V. Bagrov, K. S. Kudryashova, O. S. Sokolova and K. V. Shaitan, J. Biomech., DOI:10.1016/j.jbiomech.2013.01.022.

15 F. Rico, P. Roca-Cusachs, N. Gavara, R. Farré, M. Rotger and D. Navajas, Phys. Rev. E, 2005, 72, 21914.

16 M. Lekka, D. Gil, K. Pogoda, J. Dulińska-Litewka, R. Jach, J. Gostek, O. Klymenko, S. Prauzner-Bechcicki, Z. Stachura, J. Wiltowska-Zuber, K. Okoń and P. Laidler, Arch Biochem Biophys, DOI:10.1016/j.abb.2011.12.013.

17 S. Kessel, S. Schmidt, R. Müller, E. Wischerhoff, A. Laschewsky, J.-F. Lutz, K. Uhlig, A. Lankenau, C. Duschl and A. Fery, Langmuir, 2010, 26, 3462–3467.

18 K. D. Jandt, M. Finke and P. Cacciafesta, Colloids Surfaces B Biointerfaces, 2000, 19, 301–314.

19 V. Vyas, A. Podestà and P. Milani, J. Nanosci. Nanotechnol., 2011, 11, 4739–4748.

20 A. Keaton, J. F. Holzrichter, R. Balhorn and W. J. Siekhaus, eds. H. J. Güntherodt, D. Anselmetti and E. Meyer, Springer, Netherlands, Dordrecht, 1995, pp. 91–97.

21 V. Vyas, N. Nagarajan, P. Zorlutuna and B. D. Huey, J. Bionanoscience, 2017, 11, 319–322.

22 I. Sokolov, MRS Bull., 2012, 37, 522–527.

23 I. D. Medalsy and D. J. Müller, ACS Nano, 2013, 7, 2642–2650.

24 S. E. Cross, Y.-S. Jin, J. Rao and J. K. Gimzewski, Nat. Nanotechnol., 2007, 2, 780–783.

25 Y. F. Dufrene, D. Martinez-Martin, I. Medalsy, D. Alsteens and D. J. Muller, Nat Meth, 2013, 10, 847–854.

26 M. Plodinec, 2012.

27 J. E. Sader, I. Larson, P. Mulvaney and L. R. White, Rev. Sci. Instrum., 1995, 66, 3789–3798.

28 N. Guz, M. Dokukin, V. Kalaparthi and I. Sokolov, Biophys. J., 2014, 107, 564–575.

29 M. E. Dokukin and I. Sokolov, Macromolecules, 2012, 45, 4277–4288.

30 J. H. Hoh and C. Schoenenberger, J. Cell Sci.

31 E.-Y. Kwon, Y.-T. Kim and D.-E. Kim, J. Mech. Sci. Technol., 2009, 23, 1932–1938.

32 K. L. Bagnoli, University of Connecticut, 2007.

33 K. Gelse, E. Pöschl and T. Aigner, Adv. Drug Deliv. Rev., 2003.

34 W. S. Leong, C. Y. Tay, H. Yu, A. Li, S. C. Wu, D. H. Duc, C. T. Lim and L. P. Tan, Biochem. Biophys. Res. Commun., DOI:10.1016/j.bbrc.2010.09.052.

35 J. P. Winer, S. Oake and P. A. Janmey, PLoS One, DOI:10.1371/journal.pone.0006382.

36 Y. L. Yang, L. M. Leone and L. J. Kaufman, Biophys. J., DOI:10.1016/j.bpj.2009.07.035.

37 A. J. Heim, W. G. Matthews and T. J. Koob, Appl. Phys. Lett., DOI:10.1063/1.2367660.

38 X. Ma, M. E. Schickel, M. D. Stevenson, A. L. Sarang-Sieminski, K. J. Gooch, S. N. Ghadiali and R. T. Hart, Biophys. J., 2013, 104, 1410–1418.

39 M. P. Wenger, L. Bozec, M. A. Horton and P. Mesquida, Biophys J, DOI:S0006-3495(07)71383-8[pii]\n10.1529/biophysj.106.103192.

40 S. Bancelin, C. Aimé, I. Gusachenko, L. Kowalczuk, G. Latour, T. Coradin and M.-C. Schanne-Klein, Nat. Commun., 2014, 5, 4920.

41 T. Ushiki, Arch. Histol. Cytol., 2002, 65, 109–126.

42 B. A. Roeder, K. Kokini, J. E. Sturgis, J. P. Robinson and S. L. Voytik-Harbin, J. Biomech. Eng., DOI:10.1115/1.1449904.

43 P. D. Yurchenco and B. L. Patton, Curr. Pharm. Des.

44 P. D. Yurchenco and J. C. Schittny, FASEB J.

45 J. Takagi, Y. Yang, J. Liu, J. Wang and T. A. Springer, Nature, DOI:10.1038/nature01873.

46 R. Kalluri, Nat. Rev. Cancer, DOI:10.1038/nrc1094.

47 E. Pöschl, U. Schlötzer-Schrehardt, B. Brachvogel, K. Saito, Y. Ninomiya and U. Mayer, Development, 2004, 131, 1619 LP–1628.

48 T. T. Goodman, P. L. Olive and S. H. Pun, Int. J. Nanomedicine.

49 B. Glimelius, B. Norling, T. Nederman and J. Carlsson, Apmis.

50 M. J. Paszek, N. Zahir, K. R. Johnson, J. N. Lakins, G. I. Rozenberg, A. Gefen, C. A. Reinhart-King, S. S. Margulies, M. Dembo, D. Boettiger, D. A. Hammer and V. M. Weaver, Cancer Cell, 2005, 8, 241–254.

51 P. P. Provenzano, D. R. Inman, K. W. Eliceiri and P. J. Keely, Oncogene, 2009, 28, 4326–4343.

52 O. Chaudhuri, S. T. Koshy, C. Branco da Cunha, J.-W. Shin, C. S. Verbeke, K. H. Allison and D. J. Mooney, Nat Mater, 2014, 13, 970–978.

53 R. Timpl, H. Wiedemann, V. Van Delden, H. Furthmayr and K. KÜhn, Eur. J. Biochem., 1981, 120, 203–211.

54 K. Kühn, H. Wiedemann, R. Timpl, J. Risteli, H. Dieringer, T. Voss and R. W. Glanville, FEBS Lett., 1981, 125, 123–128.

55 M. W. Tibbitt and K. S. Anseth, Biotechnol. Bioeng., 2009.

56 A. Sawhney, C. Pathak and J. Hubbell, Macromolecules, DOI:10.1021/ma00056a005.

57 M. P. Lutolf, Integr. Biol., DOI:10.1039/b902243k.

58 F. Pampaloni, E. G. Reynaud and E. H. K. Stelzer, Nat. Rev. Mol. Cell Biol., 2007, 8, 839–845.

